# Discovery of a Novel Inhibitor for Chikungunya Virus

**DOI:** 10.1101/2022.03.23.485574

**Authors:** Ana C. Puhl, Rafaela S. Fernandes, Andre S. Godoy, Laura H. V. G. Gil, Glaucius Oliva, Sean Ekins

**Author notes:** Both authors contribute equally: Ana C. Puhl, Rafaela S. Fernandes.

## Abstract

Chikungunya virus (CHIKV) is the etiological agent of chikungunya fever, a (re)emerging arbovirus infection, that causes severe and often persistent arthritis, representing a serious health problem worldwide for which no antivirals are currently available. Despite the efforts over the last decade to identify and optimize new inhibitors or to reposition existing drugs, no compound has progressed to clinical trials and prophylaxis is based on vector control, which has shown limited success in containing the virus. Herein, we screened 36 compounds using a replicon system and ultimately identified 3-methyltoxoflavin with activity against CHIKV using cell assays (EC_50_ 200 nM). We have additionally screened 3-methyltoxoflavin against a panel of viruses and showed that it also inhibits yellow fever virus (IC_50_ 370 nM, SI=3.2 in Huh-7 cells). 3-methyltoxoflavin is a known protein disulfide isomerase (PDI) inhibitor, and inhibitor of alphaviruses which likely depends on this host protein to aid in facilitating disulfide bond formation and isomerization, since alphaviruses require conserved cysteine residues for proper folding and assembly of the E1 and E2 envelope glycoproteins. In summary we demonstrate that 3-methyltoxoflavin has activity against CHIKV and may represent a starting point for optimization to develop inhibitors to this and other viruses.

## Introduction

Chikungunya virus (CHIKV), a member of the *Alphavirus* genus (*Togaviridae* family), is the etiologic agent of Chikungunya fever, a globally spreading mosquito-borne disease, first isolated in 1952, which causes periodic and explosive outbreaks of a severe and often persistent arthritis ^1^. CHIKV has reemerged as a significant public health threat since the 2005 in La Réunion. In February 2005, a major outbreak occurred on the islands of the Indian Ocean, followed by a large number of CHIKV infection cases reported in Europe and India in 2006 and 2007, respectively ^2^. In December 2013, CHIKV emerged on the American continent, on the island of St. Martin, Caribbean, and by the end of December 2015, nearly 1 million cases had been notified in the Americas ^3, 4^. The reemergence of the virus in many parts of the world represents a serious public health concern ^5^. Since 2013, hundreds of thousands of cases of the disease were confirmed in the north and northwest parts of Brazil ^6^. The outbreak is still not contained, and the risk of expansion to the southern part of Brazil and other countries still exists.

Driven by the medical importance of this virus, as well as the lack of approved antivirals, research into the field of CHIKV antivirals has recently intensified. CHIKV is transmitted to humans by *Aedes sp.* mosquitoes, mainly *Aedes aegypti* and *A. albopictus*, both widely disseminated in tropical and subtropical regions ^7^. Humans are the primary host of CHIKV during epidemic periods. Rare *in utero* transmission has been documented mostly during the second trimester. Intrapartum transmission has also been documented when the mother was viremic around the time of delivery ^8^. Most people infected with CHIKV become symptomatic. The incubation period is typically 3–7 days (range, 1–12 days).

The viral infection by CHIKV in humans is distinguished by a very high viremia and debilitating symptoms such as joint pain and inflammation, fever, rash and even morbidity that may persist for years ^9, 10^. To date there are no drug treatments or vaccines available to contain the infection. Without specific vaccines or drugs against CHIKV, prophylaxis is based on vector control, which has shown limited success in containing the virus ^9, 11^. Treatment for symptoms can include rest, fluids, and use of non-steroidal anti-inflammatory drugs (NSAIDs) to relieve acute pain and fever. Therefore, effective treatments against CHIKV would mitigate viral spread and limit morbidity associated with the disease ^10^. Owing to mechanisms that are poorly understood, recurrent and persistent myalgia and arthralgia have been reported to last for years after the infection clears in some patients. A recent study with a macaque model suggested that the chronic phase could be caused by inflammatory responses toward persistent CHIKV in certain tissues, rather than an autoimmune-mediated response, as was initially believed^12^.

Some of the well-known broad-spectrum antivirals like ribavirin and interferon, may prove to be promising against CHIKV^13^ although there is no evidence supporting the clinical efficacy they could be subject to clinical trials in future^13, 14^. Recent drug discovery efforts have included computer-aided design to identify an inhibitor of the E2-E1 envelope glycoprotein complex leading to an optimal compound with and EC_50_ of 1.6 mM ^15^. Several classes of CHIKV inhibitors have been described such as the benzoannulene replication inhibitors ^16^, pyrimidones ^17^, indoles ^18^ and others ^19^ derived by fragment-based screens or high throughput screening. Despite the efforts performed in the last decade to identify and optimize new inhibitors or to reposition existing drugs, no compound has progressed towards clinical trials.

## Methods

### Cells and compounds

BHK-21 [C-13] and HepG2 cell lines were purchased from Banco de Células do Rio de Janeiro (BCRJ) and maintained in Dulbecco’s Modified Eagle’s Medium (DMEM, GIBCO) containing 10% heat-inactivated fetal bovine serum (FBS, GIBCO), 100 units/ml penicillin and 100 μg/ml streptomycin at 37 °C in a 5% CO_2_-humidified incubator. BHK-21-GLuc-nSP-CHIKV-99659 cell line, harboring a Chikungunya virus replicon expressing Gaussia luciferase (Gluc) and neomycin phosphotransferase (Neo) genes, were maintained in DMEM 10% FBS with 500 μg/ml G418 (Sigma-Aldrich). The development and characterization of this CHIKV replicon cell line will be described elsewhere. CPI compounds (> 90% purity) were solubilized in 100% DMSO (v/v) and further diluted with assay media to a final DMSO concentration of 1% (v/v) for the antiviral assays.

### Replicon-based antiviral assays

A total of 36 compounds were evaluated as potential inhibitors of the viral replication using the BHK-21-T7-Gluc-nSP-CHIKV-99659 cell line. Primary screening of each compound was performed at 20 μM with 1% DMSO final concentration in a 96-well format. Approximately 2 × 10^4^ replicon cells/well in DMEM 10% FBS were seeded in a 96-well plate. After 16 h of incubation at 37 °C 5% CO_2_, medium was replaced with fresh DMEM supplemented with 2% FBS and compounds were added to the cells. After a 48 h-incubation, 40 μL of the cells’ supernatant containing secreted GLuc were mixed with *Renilla* luciferase Assay Reagent (100 μl) (Promega). GLuc activity was measured using SpectraMax i3 Multi-mode Detection Platform (Molecular Devices).

Compounds that inhibited GLuc activity in ≥ 80% were assayed for the determination of their effective (EC_50_) and cytotoxicity (CC_50_) concentrations. As described above, replicon cells were seeded in 96-well plates and after 16 h of incubation at 37 °C 5% CO_2_, medium was replaced with fresh DMEM with 2% FBS and compounds at 2-fold serial dilutions were added to the cells. After a 48 h-incubation, 40 μL of the cells’ supernatant were mixed with *Renilla* luciferase Assay Reagent (100 μL) (Promega) and Gluc activity were measured using SpectraMax i3 Multi-mode Detection Platform (Molecular Devices). The compounds concentrations required to inhibit 50% of the Gluc activity (EC_50_) were calculated using the OriginPro 9.0 software. BHK-21 cells in 1% DMSO were used as negative control.

The cytotoxicity of compounds were evaluated by a cell proliferation-based MTT (3-(4,5-dimethylthiazol-2-yl)-2,5-diphenyltetrazolium bromide) assay^20^. Replicon cells were seeded in 96-well plates as described above. After 48 h of incubation with the compounds at 2-fold serial dilutions, MTT (3-(4,5-dimethylthiazol-2-yl)-2,5-diphenyltetrazolium bromide) solution (5mg/mL in PBS) was added to the wells at one tenth of the well volume and the plates were incubated at 37 ºC 5% CO_2_ for 3-4 hours. Next, the medium was removed, and formazan crystals were solubilized in DMSO. Absorbance was measured at 570 nm wavelength in SpectraMax 384 plate reader (Molecular Devices). The compounds concentrations required to cause 50% cytotoxicity (CC_50_) were estimated using the OriginPro 9.0 software. All the antiviral assays were performed twice in duplicates.

### Cytotoxicity of compound 3-methyltoxoflavin in HepG2 cells

The cell proliferation-based MTT was also used to evaluate the potential drug-induced hepatotoxicity of compound 3-methyltoxoflavin using hepatocellular carcinoma cells (HepG2). Approximately 2 × 10^4^ HepG2 cells per well were seeded in 96-well plates and the assay was performed as described above. The compound concentration required to cause 50% cytotoxicity (CC_50_) was estimated using the OriginPro 9.0 software. The assay was performed twice in duplicates.

### Primary CPE and secondary VYR assays for viruses

We screened the following viruses: Dengue virus 2, Eastern equine encephalitis virus, Enterovirus-71, MERS-Coronavirus, Influenza A (H1N1), Japanese encephalitis virus, Mayaro virus, Measles, Respiratory syncytial virus, Rift Valley fever virus, Tacaribe virus, Usutu Virus, Venezuelan equine encephalitis virus, West Nile virus, Western Equine Encephalitis and Zika Virus (Table S1). The assays were conducted as described ^21^. Two sets of data are provided: (1) EC_50_, CC_50_, and SI_50_ obtained from the neutral red assay, and (2) EC_90_ (obtained from the VYR assay), CC_50_ (same value as for item 1), and SI_90_.

## Data analysis

For the replicon-based assays, statistical calculations of Z′-values were made as follows: Z′ =1– ((3SD of sample + 3SD of control)/|Mean of sample – Mean of control|). Here, SD is the standard deviation of the luminescent signals from cell control (BHK-21) or sample. Z′ values between 0.5 and 1 are considered good quality^20, 22^.

## Results and Discussion

CHIKV is a (re)emerging arbovirus and about 3–5 million cases of CHIKV infections are reported every year globally (World Health and Organization (WHO), 2021. After an acute phase with fever and joint arthralgia, viral infection can lead to chronic debilitating arthritis, which has important social and economic consequences. Although mortality is low, in neonates, elderly people or patients suffering from diabetes or heart disease, infection with CHIKV can lead to severe complications including death ^23^. On the other hand, morbidity is high, with important social and economic consequences. Moreover, coinfections with dengue virus, transmitted by the same vector, also raises serious concerns ^24^.

We screened 36 compounds (Table 1) using BHK-21-GLuc-nSP-CHIKV-99659 replicon cell line. Each compound was first assayed at 20 μM concentration in a 96-well format plate (Table 1). Nine compounds inhibited the luciferase activity (furaltadone, amodiaquine, promazine, fluphenazine, quinacrine, 4-hydroxyderricin, xanthoangelol, (2E)-1-[3-(2-Hydroperoxy-3-methyl-3-buten-1-yl)-2-hydroxy-4-methoxyphenyl]-3-(4-hydroxyphenyl)-2-propen-1-one, 3-methyltoxoflavin) in ≥ 80% and were further assayed for the determination of their EC_50_ and CC_50_ values. These compounds were tested in a dose-dependent manner, and only xanthoangelol, quinacrine and 3-methyltoxoflavin showed activity. Xanthoangelol and quinacrine decreased the GLuc signals in a dose-dependent manner, with EC_50_ values in the low micromolar range (Figure 1 A,B). Those compounds exhibited high cytotoxicity to the replicon cells, resulting in poor selectivity indexes (CC_50_/EC_50_, SI)) of 1.66 and 1.64, respectively (Figure 1A,B). In contrast, 3-methyltoxoflavin exhibited a low cytotoxicity to the cells with an EC_50_ of 20 ± 0.04 μM and SI=30.8 (Figure 1C). We also evaluated 3-methyltoxoflavin hepatotoxicity using HepG2 cells, which resulted in a CC_50_ of 11.0 ± 1.73 μM (Figure 1D), showing that this compound displays a moderate to low toxicity to human liver cells. 3-methyltoxoflavin is a protein disulfide isomerase (PDI) inhibitor^25^.

**Table 1:**
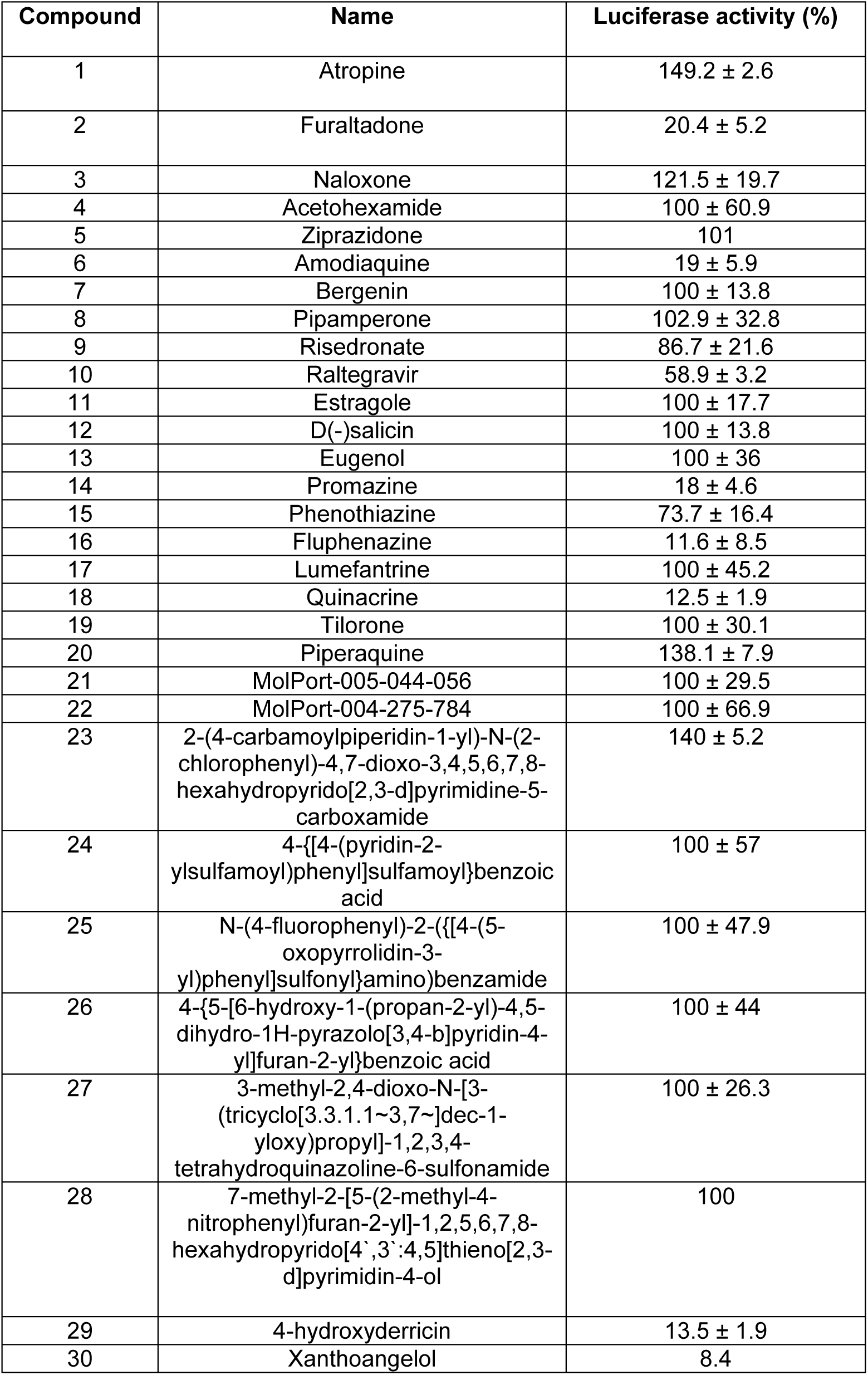
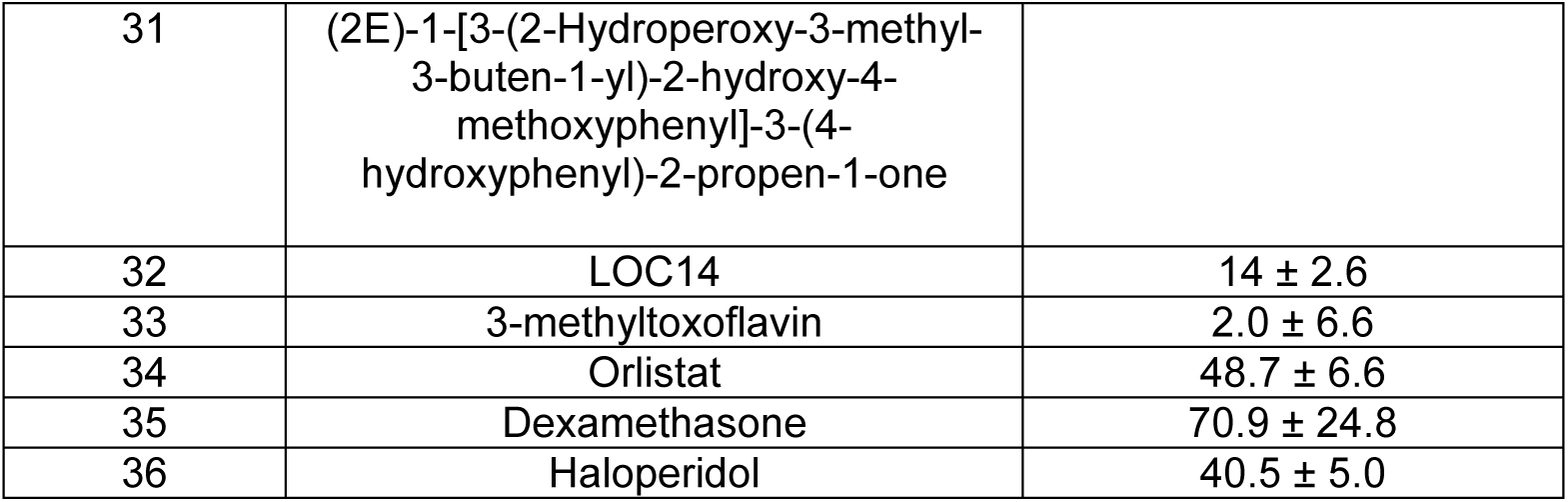
Antiviral screening of CPI compounds at 20 μM using the CHIKV reporter replicon cell line BHK-21-Gluc-nSP-CHIKV-99659. Average results and standard deviations of two independent assays performed in duplicates.

**Figure 1.**
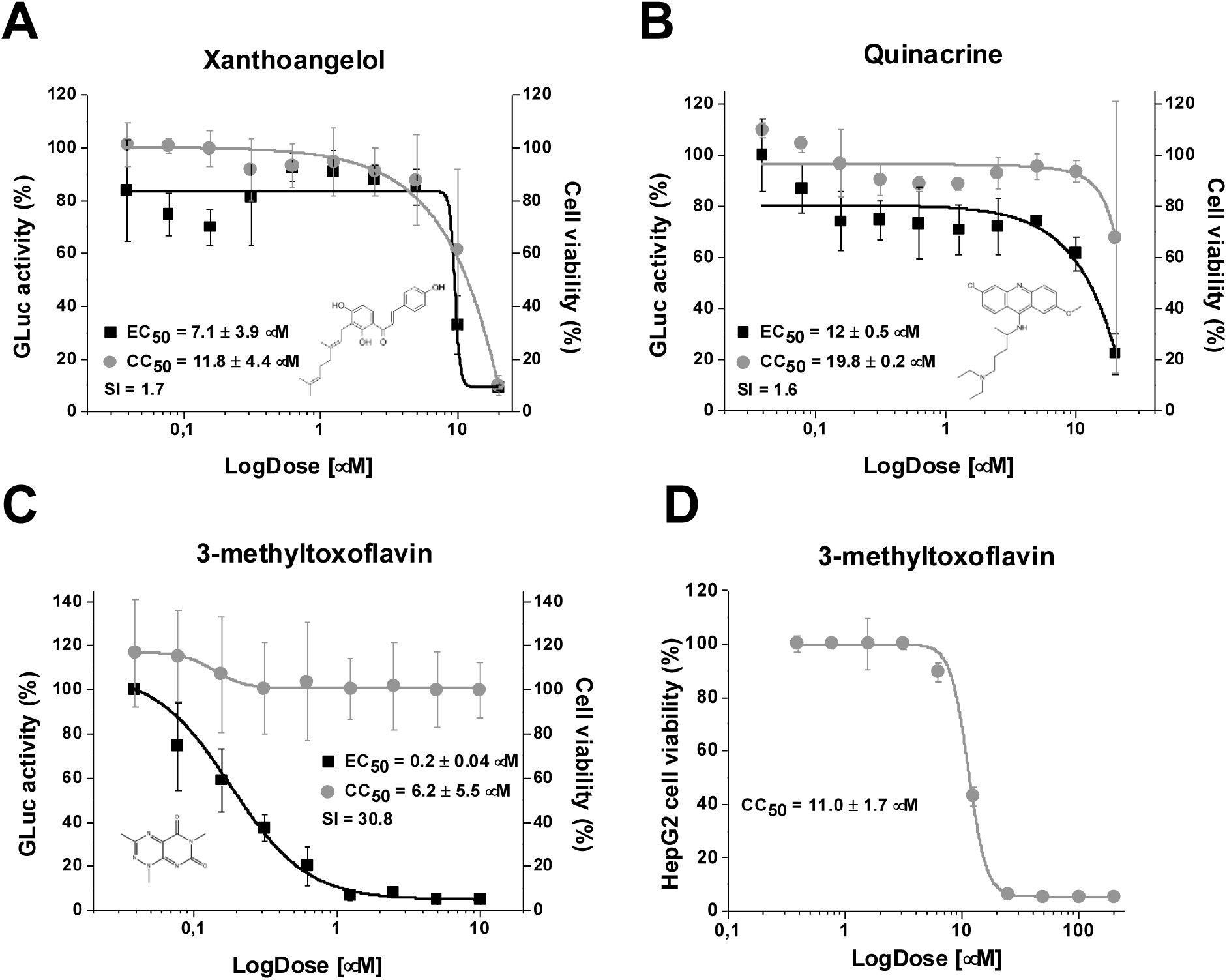
The dose-response curves. **A)** Xanthoangelol, B) Quinacrine and **3)** 3-methyltoxoflavin tested in replicon cell lines. CHIKV replicon cells were incubated with compounds at 2-fold serial dilutions (from 20 μM to 0.03 μM) for 48 h and Gluc activity was measured from cells’ supernatant. **D)** Cytotoxicity of 3-methyltoxoflavin was tested in HepG2 cells. The cells were incubated with compound at 2-fold serial dilutions (from 200 μM to 0.39 μM) for 48 h and MTT (3-(4,5-dimethylthiazol-2-yl)-2,5-diphenyltetrazolium bromide) solution (5 mg/mL in PBS) were added to the wells. After 3-4 h of incubation, the formazan crystals were solubilized in DMSO, and absorbance was read at 570 nm. Average results of two independent experiments. Error bars represents the standard deviations.

Alphaviruses require conserved cysteine residues for proper folding and assembly of the E1 and E2 envelope glycoproteins^26^, and likely depends on host PDI to aid in facilitating disulfide bond formation and isomerization in these proteins. 3-Methyltoxoflavin is a potent PDI inhibitor, with an IC_50_ of 170 nM ^25^ and we hypothesize that the mechanism for the activity of 3-methyltoxoflavin may be due in part to inhibition of PDI ^25^. PDI is a promising target for cancer therapy ^25, 27^ and many viruses, such as HIV ^28^, CHIKV ^29^, Dengue ^30, 31^, Zika ^31^, Hepatitis C ^32^, influenza ^33^, as well as others ^34^. 3-Methyltoxoflavin also induces Nrf2 antioxidant response, ER stress response, and autophagy^25^. 3-Methyltoxoflavin also bears some 2D structural resemblance to nucleotide and nucleoside type inhibitors so this may also represent another mechanism by interfering directly with viral targets.

So far, a few compounds have been described with activity for CHIKV(^15,16,17,18,19^). Previous work performed using a CHIKV replicon cell line, screened ∼3000 bioactive molecules including FDA approved drugs and identified the isoquinoline alkaloid berberine as a potent inhibitor of CHIKV replication ^35^. The efficacy of this compound was tested in a CHIKV-induced arthritis mouse model and showed the alleviation of the inflammatory symptoms ^36^. Another approach has been described to identify effective antiviral drugs against CHIKV based on a human genome-wide loss-of-function screen ^37^, which resulted in identification of important targets and the combination of two existing drugs, 5-(tetradecyloxy)-2-furoic acid (TOFA, and pimozide, targeting fatty acid synthesis and calmodulin signaling respectively, that showed anti-CHIKV activity *in vitro* and *in vivo* ^37^.

We have demonstrated that 3-methyltoxoflavin has activity against CHIKV and was also screened a panel of viruses (Table S1), and only showed activity against yellow fever (YFV 17D strain) with an EC_50_ 0.37 μM and SI 3.2 in the visual cytopathic toxicity assay (Table 2). CHIKV and YFV are both arboviruses transmitted by the *Aedes aegypti* mosquito, however CHIKV belongs to the family *Togaviridae* and YFV belongs to the *Flaviviridae*. The mechanism of action remains to be fully elucidated but it may suggest that 3-methyltoxoflavin could be further optimized to develop novel therapeutic agents to treat these viruses.

**Table 2:** 3-methyltoxoflavin was tested against CHIKV and Yellow Fever.

| **Virus/strain** | **Cell type** | **Drug Assay Name** | **EC<sub>50</sub>**<br>**(µM)** | **EC<sub>90</sub>**<br>**(µM)** | **CC<sub>50</sub>**<br>**(µM)** | **SI<sub>50</sub>**<br>**(µM)** | **SI<sub>90</sub>**<br>**(µM)** |
| --- | --- | --- | --- | --- | --- | --- | --- |
| Chikungunya virus/ S27 | Huh-7 | Visual (cytopathic effect/toxicity) | 0.19 |  | 3.2 | 17 |  |
| Chikungunya virus/ S27 | Huh-7 | Neutral Red (cytopathic effect/toxicity) | <0.1 |  | 1.1 | >11 |  |
| Chikungunya virus/ S27 | Vero 76 | Visual (Virus yield reduction)/Neutral Red (Toxicity) |  | >1.7 | 1.7 |  | 0 |
| Chikungunya virus/ S27 | Vero 76 | Neutral Red (Cytopathic effect/Toxicity) | >1.7 |  | 1.7 | 0 |  |
| Yellow Fever Virus/ YFV 17D | Huh-7 | Visual (cytopathic effect/toxicity) | 0.37 |  | 1.2 | 3.2 |  |
| Yellow Fever Virus/ YFV 17D | Huh-7 | Neutral Red (cytopathic effect/toxicity) | 0.42 |  | 1.2 | 2.9 |  |

## Acknowledgments

We kindly acknowledge Dr. Mindy Davis and colleagues for assistance with antiviral testing services through NIAID and our colleagues at Collaborations Pharmaceuticals, Inc. for their assistance in the early stages of this project. We kindly acknowledge Dr. Scott M. Laster and Dr. Nadja Cech for providing xanthoangelol.

We kindly acknowledge NIH funding: R44GM122196-02A1 from NIGMS (PI – Sean Ekins),

Collaborations Pharmaceuticals, Inc. has utilized the non-clinical and pre-clinical services program offered by the National Institute of Allergy and Infectious Diseases.

LHVGG would like to thank CNPq (grant 440773/2019-8).

RSF and GO received financial support from Fundação de Amparo à Pesquisa do Estado de São Paulo (FAPESP), grant 2018/05130-3 to RSF and CEPID grant 2013/07600-3 to GO.

## Supplemental Material

**Table S1:**
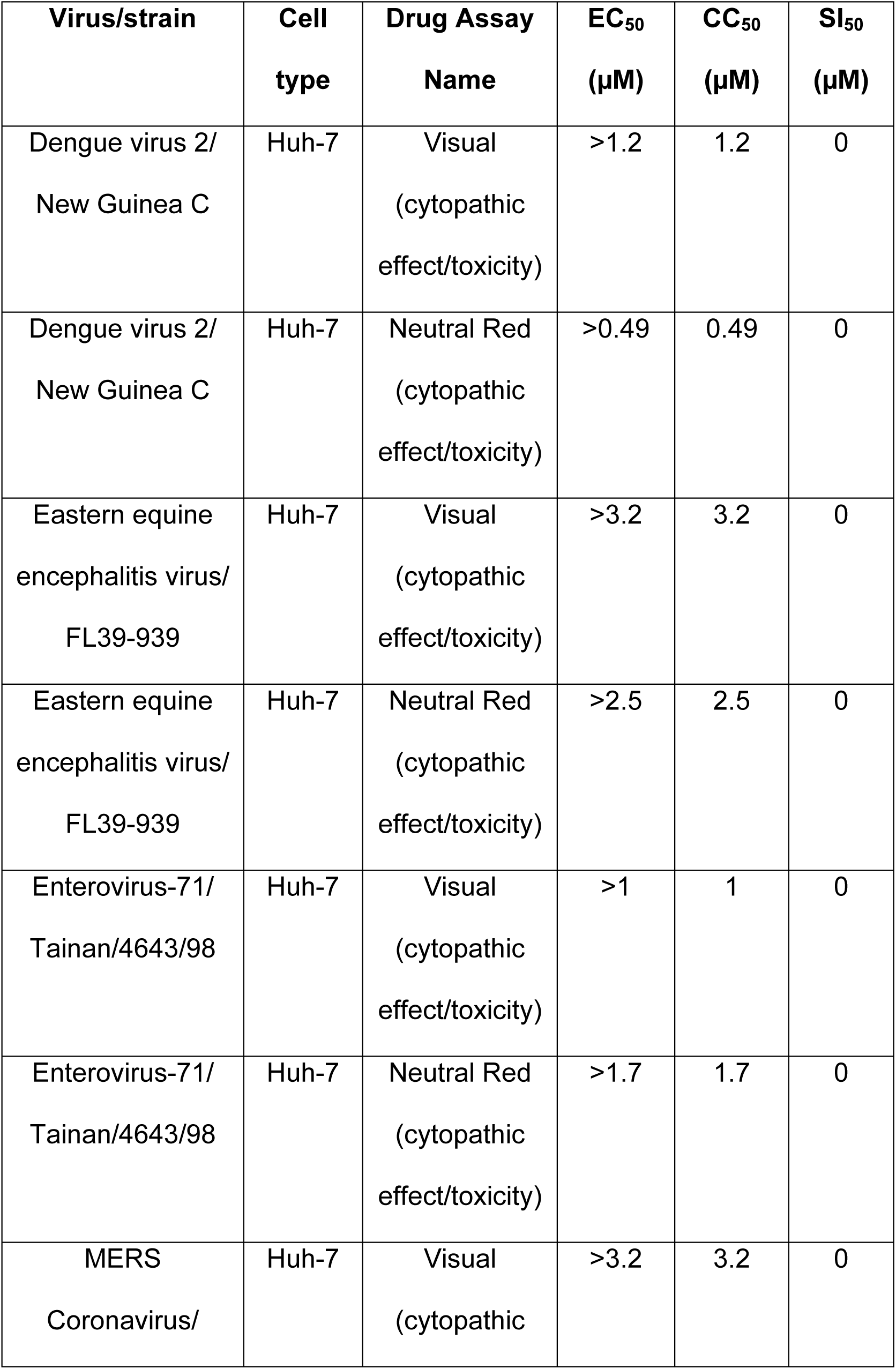
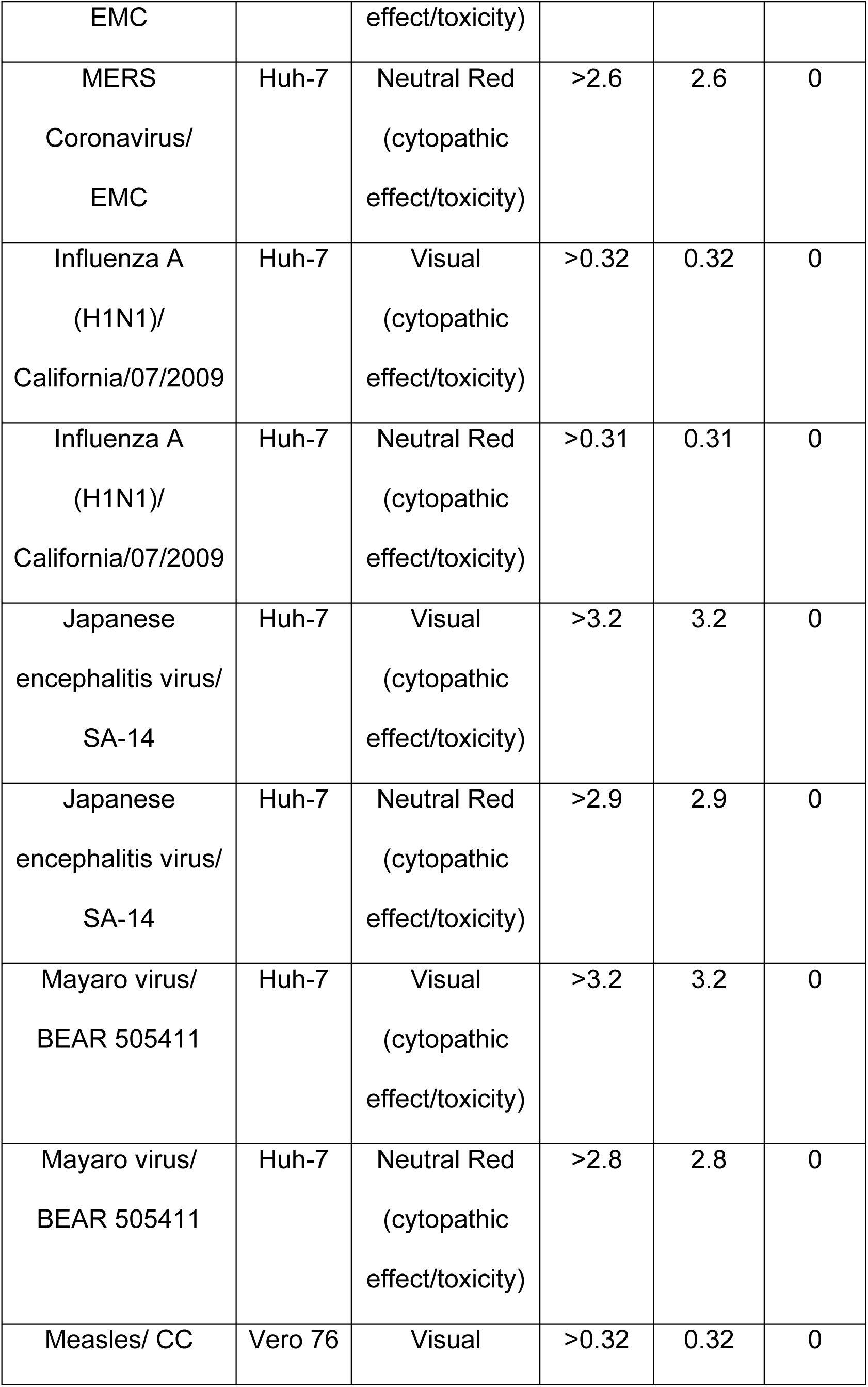
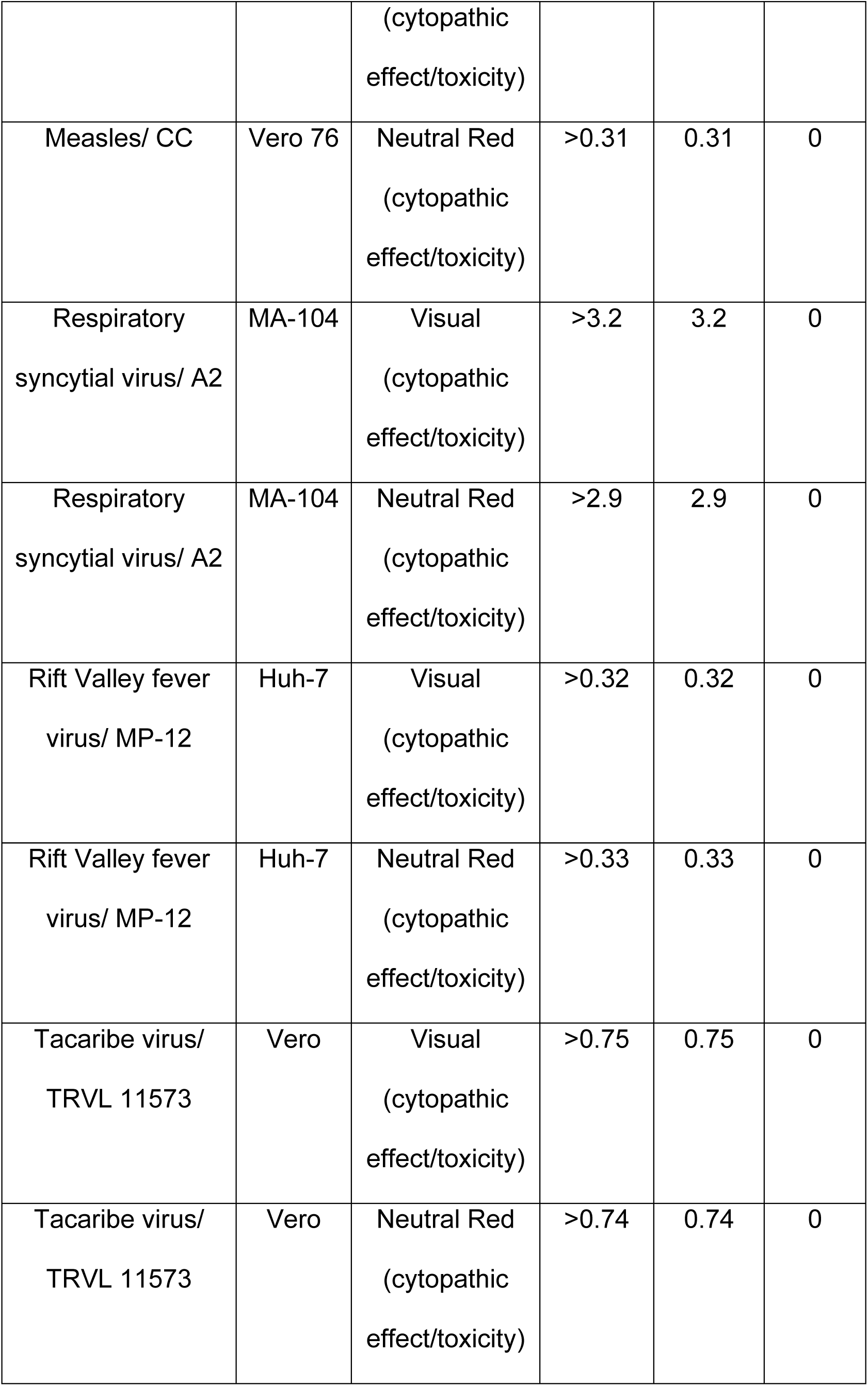
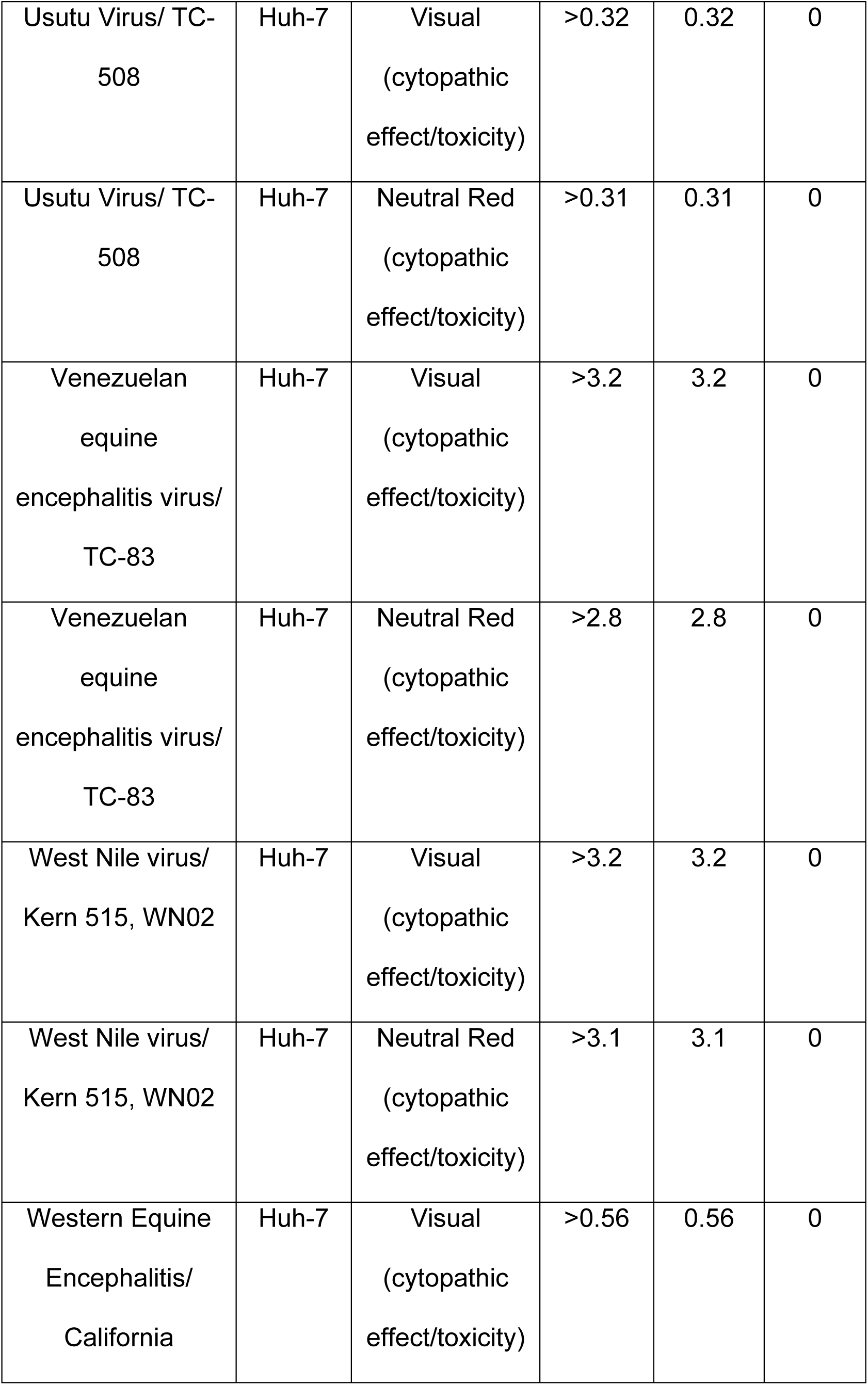
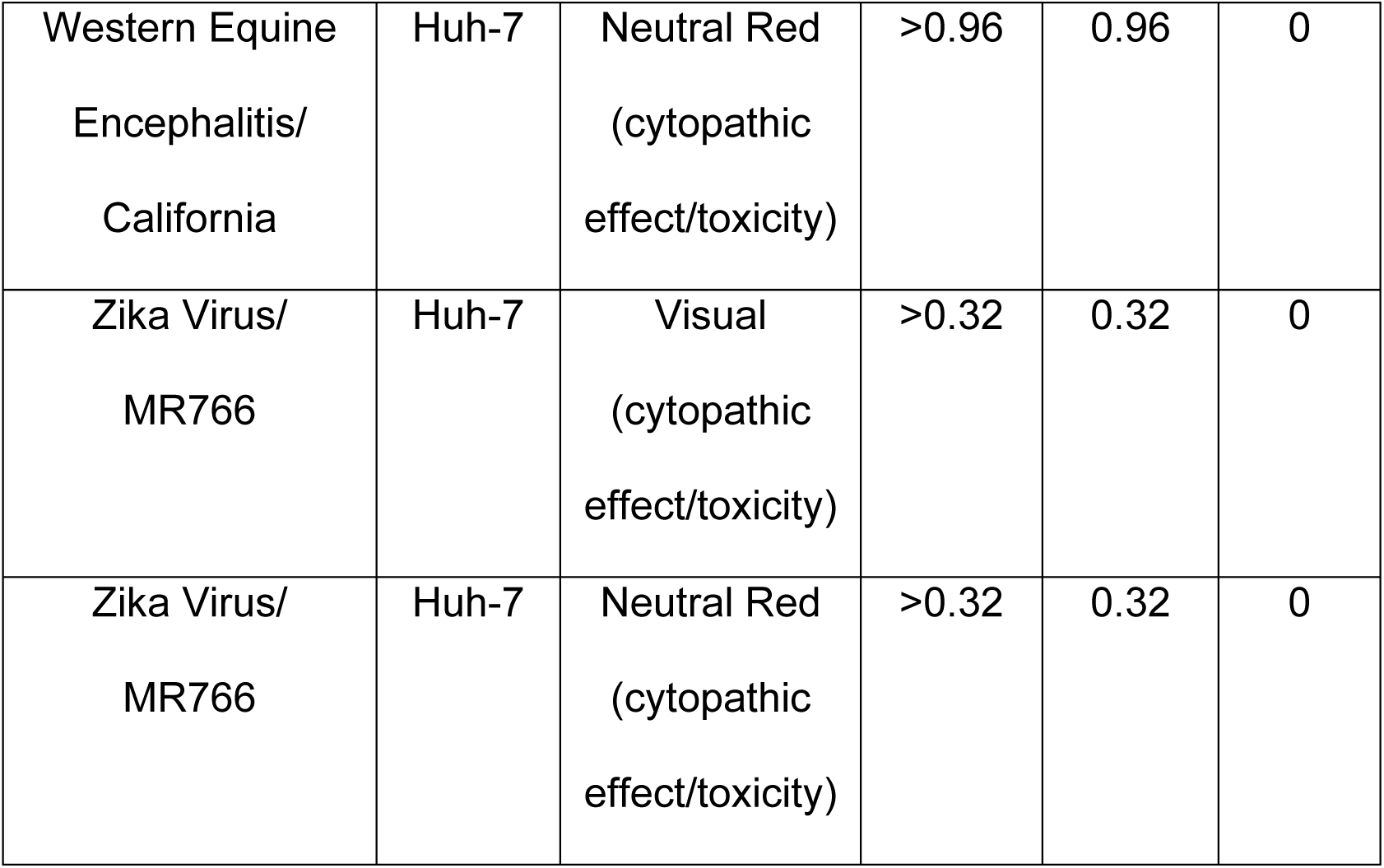
3-methyltoxoflavin was tested against several viruses.

## References

(1) Mason, P. J.; Haddow, A. J. An epidemic of virus disease in Southern Province, Tanganyika Territory, in 1952-53; an additional note on Chikungunya virus isolations and serum antibodies. Trans R Soc Trop Med Hyg 1957, 51 (3), 238–240. DOI: 10.1016/0035-9203(57)90022-6.

(2) Schuffenecker, I.; Iteman, I.; Michault, A.; Murri, S.; Frangeul, L.; Vaney, M. C.; Lavenir, R.; Pardigon, N.; Reynes, J. M.; Pettinelli, F.; et al. Genome microevolution of chikungunya viruses causing the Indian Ocean outbreak. PLoS Med 2006, 3 (7), e263. DOI: 10.1371/journal.pmed.0030263. Panning, M.; Grywna, K.; van Esbroeck, M.; Emmerich, P.; Drosten, C. Chikungunya fever in travelers returning to Europe from the Indian Ocean region, 2006. Emerg Infect Dis 2008, 14 (3), 416–422. DOI: 10.3201/eid1403.070906.

(3) Wahid, B.; Ali, A.; Rafique, S.; Idrees, M. Global expansion of chikungunya virus: mapping the 64-year history. Int J Infect Dis 2017, 58, 69–76. DOI: 10.1016/j.ijid.2017.03.006.

(4) Rodrigues Faria, N.; Lourenço, J.; Marques de Cerqueira, E.; Maia de Lima, M.; Pybus, O.; Carlos Junior Alcantara, L. Epidemiology of Chikungunya Virus in Bahia, Brazil, 2014-2015. PLoS Curr 2016, 8. DOI: 10.1371/currents.outbreaks.c97507e3e48efb946401755d468c28b2.

(5) Silva, J. V. J.; Ludwig-Begall, L. F.; Oliveira-Filho, E. F.; Oliveira, R. A. S.; Durães-Carvalho, R.; Lopes, T. R. R.; Silva, D. E. A.; Gil, L. H. V. G. A scoping review of Chikungunya virus infection: epidemiology, clinical characteristics, viral co-circulation complications, and control. Acta Trop 2018, 188, 213–224. DOI: 10.1016/j.actatropica.2018.09.003.

(6) Aguiar, B. S.; Lorenz, C.; Virginio, F.; Suesdek, L.; Chiaravalloti-Neto, F. Potential risks of Zika and chikungunya outbreaks in Brazil: A modeling study. Int J Infect Dis 2018, 70, 20–29. DOI: 10.1016/j.ijid.2018.02.007.

(7) Coffey, L. L.; Failloux, A. B.; Weaver, S. C. Chikungunya virus-vector interactions. Viruses 2014, 6 (11), 4628–4663. DOI: 10.3390/v6114628.

(8) Charlier, C.; Beaudoin, M. C.; Couderc, T.; Lortholary, O.; Lecuit, M. Arboviruses and pregnancy: maternal, fetal, and neonatal effects. Lancet Child Adolesc Health 2017, 1 (2), 134–146. DOI: 10.1016/S2352-4642(17)30021-4.

(9) Burt, F. J.; Chen, W.; Miner, J. J.; Lenschow, D. J.; Merits, A.; Schnettler, E.; Kohl, A.; Rudd, P. A.; Taylor, A.; Herrero, L. J.; et al. Chikungunya virus: an update on the biology and pathogenesis of this emerging pathogen. Lancet Infect Dis 2017, 17 (4), e107–e117. DOI: 10.1016/S1473-3099(16)30385-1.

(10) Silva, L. A.; Dermody, T. S. Chikungunya virus: epidemiology, replication, disease mechanisms, and prospective intervention strategies. J Clin Invest 2017, 127 (3), 737–749. DOI: 10.1172/JCI84417.

(11) Vairo, F.; Haider, N.; Kock, R.; Ntoumi, F.; Ippolito, G.; Zumla, A. Chikungunya: Epidemiology, Pathogenesis, Clinical Features, Management, and Prevention. Infect Dis Clin North Am 2019, 33 (4), 1003–1025. DOI: 10.1016/j.idc.2019.08.006.

(12) Labadie, K.; Larcher, T.; Joubert, C.; Mannioui, A.; Delache, B.; Brochard, P.; Guigand, L.; Dubreil, L.; Lebon, P.; Verrier, B.; et al. Chikungunya disease in nonhuman primates involves long-term viral persistence in macrophages. J Clin Invest 2010, 120 (3), 894–906. DOI: 10.1172/JCI40104. Dupuis-Maguiraga, L.; Noret, M.; Brun, S.; Le Grand, R.; Gras, G.; Roques, P. Chikungunya disease: infection-associated markers from the acute to the chronic phase of arbovirus-induced arthralgia. PLoS Negl Trop Dis 2012, 6 (3), e1446. DOI: 10.1371/journal.pntd.0001446.

(13) Kaur, P.; Chu, J. J. Chikungunya virus: an update on antiviral development and challenges. Drug Discov Today 2013, 18 (19-20), 969–983. DOI: 10.1016/j.drudis.2013.05.002.

(14) Hucke, F. I. L.; Bugert, J. J. Current and Promising Antivirals Against Chikungunya Virus. Front Public Health 2020, 8, 618624. DOI: 10.3389/fpubh.2020.618624.

(15) Battini, L.; Fidalgo, D. M.; Alvarez, D. E.; Bollini, M. Discovery of a Potent and Selective Chikungunya Virus Envelope Protein Inhibitor through Computer-Aided Drug Design. ACS Infect Dis 2021, 7 (6), 1503–1518. DOI: 10.1021/acsinfecdis.0c00915.

(16) Ahmed, S. K.; Haese, N. N.; Cowan, J. T.; Pathak, V.; Moukha-Chafiq, O.; Smith, V. J.; Rodzinak, K. J.; Ahmad, F.; Zhang, S.; Bonin, K. M.; et al. Targeting Chikungunya Virus Replication by Benzoannulene Inhibitors. J Med Chem 2021, 64 (8), 4762–4786. DOI: 10.1021/acs.jmedchem.0c02183.

(17) Zhang, S.; Garzan, A.; Haese, N.; Bostwick, R.; Martinez-Gzegozewska, Y.; Rasmussen, L.; Streblow, D. N.; Haise, M. T.; Pathak, A. K.; Augelli-Szafran, C. E.; et al. Pyrimidone inhibitors targeting Chikungunya Virus nsP3 macrodomain by fragment-based drug design. PLoS One 2021, 16 (1), e0245013. DOI: 10.1371/journal.pone.0245013.

(18) Scuotto, M.; Abdelnabi, R.; Collarile, S.; Schiraldi, C.; Delang, L.; Massa, A.; Ferla, S.; Brancale, A.; Leyssen, P.; Neyts, J.; et al. Discovery of novel multi-target indole-based derivatives as potent and selective inhibitors of chikungunya virus replication. Bioorg Med Chem 2017, 25 (1), 327–337. DOI: 10.1016/j.bmc.2016.10.037.

(19) Abdelnabi, R.; Kovacikova, K.; Moesslacher, J.; Donckers, K.; Battisti, V.; Leyssen, P.; Langer, T.; Puerstinger, G.; Querat, G.; Li, C.; et al. Novel Class of Chikungunya Virus Small Molecule Inhibitors That Targets the Viral Capping Machinery. Antimicrob Agents Chemother 2020, 64 (7). DOI: 10.1128/AAC.00649-20. Feibelman, K. M.; Fuller, B. P.; Li, L.; LaBarbera, D. V.; Geiss, B. J. Identification of small molecule inhibitors of the Chikungunya virus nsP1 RNA capping enzyme. Antiviral Res 2018, 154, 124–131. DOI: 10.1016/j.antiviral.2018.03.013.

(20) Li, J. Q.; Deng, C. L.; Gu, D.; Li, X.; Shi, L.; He, J.; Zhang, Q. Y.; Zhang, B.; Ye, H. Q. Development of a replicon cell line-based high throughput antiviral assay for screening inhibitors of Zika virus. Antiviral Res 2018, 150, 148–154. DOI: 10.1016/j.antiviral.2017.12.017.

(21) Puhl, A. C.; Fritch, E. J.; Lane, T. R.; Tse, L. V.; Yount, B. L.; Sacramento, C. Q.; Fintelman-Rodrigues, N.; Tavella, T. A.; Maranhão Costa, F. T.; Weston, S.; et al. Repurposing the Ebola and Marburg Virus Inhibitors Tilorone, Quinacrine, and Pyronaridine:. ACS Omega 2021, 6 (11), 7454–7468. DOI: 10.1021/acsomega.0c05996.

(22) Elshabrawy, H. A.; Fan, J.; Haddad, C. S.; Ratia, K.; Broder, C. C.; Caffrey, M.; Prabhakar, B. S. Identification of a broad-spectrum antiviral small molecule against severe acute respiratory syndrome coronavirus and Ebola, Hendra, and Nipah viruses by using a novel high-throughput screening assay. J Virol 2014, 88 (8), 4353–4365. DOI: 10.1128/JVI.03050-13. Li, Q.; Maddox, C.; Rasmussen, L.; Hobrath, J. V.; White, L. E. Assay development and high-throughput antiviral drug screening against Bluetongue virus. Antiviral Res 2009, 83 (3), 267–273. DOI: 10.1016/j.antiviral.2009.06.004.

(23) Couderc, T.; Lecuit, M. Chikungunya virus pathogenesis: From bedside to bench. Antiviral Res. 2015, 121, 120–131. DOI: 10.1016/j.antiviral.2015.07.002. Thiberville, S. D.; Moyen, N.; Dupuis-Maguiraga, L.; Nougairede, A.; Gould, E. A.; Roques, P.; de Lamballerie, X. Chikungunya fever: Epidemiology, clinical syndrome, pathogenesis and therapy. Antiviral Res. 2013, 99 (3), 345–370. DOI: 10.1016/j.antiviral.2013.06.009.

(24) Lo Presti, A.; Cella, E.; Angeletti, S.; Ciccozzi, M. Molecular epidemiology, evolution and phylogeny of Chikungunya virus: An updating review. Infect. Genet. Evol. 2016, 41, 270–278. DOI: 10.1016/j.meegid.2016.04.006Scopus.

(25) Kyani, A.; Tamura, S.; Yang, S.; Shergalis, A.; Samanta, S.; Kuang, Y.; Ljungman, M.; Neamati, N. Discovery and Mechanistic Elucidation of a Class of Protein Disulfide Isomerase Inhibitors for the Treatment of Glioblastoma. ChemMedChem 2018, 13 (2), 164–177. DOI: 10.1002/cmdc.201700629.

(26) Voss, J. E.; Vaney, M. C.; Duquerroy, S.; Vonrhein, C.; Girard-Blanc, C.; Crublet, E.; Thompson, A.; Bricogne, G.; Rey, F. A. Glycoprotein organization of Chikungunya virus particles revealed by X-ray crystallography. Nature 2010, 468 (7324), 709–712. DOI: 10.1038/nature09555.

(27) Xu, S.; Sankar, S.; Neamati, N. Protein disulfide isomerase: a promising target for cancer therapy. Drug Discov Today 2014, 19 (3), 222–240. DOI: 10.1016/j.drudis.2013.10.017.

(28) Khan, M. M.; Simizu, S.; Lai, N. S.; Kawatani, M.; Shimizu, T.; Osada, H. Discovery of a small molecule PDI inhibitor that inhibits reduction of HIV-1 envelope glycoprotein gp120. ACS Chem Biol 2011, 6 (3), 245–251. DOI: 10.1021/cb100387r.

(29) Langsjoen, R. M.; Auguste, A. J.; Rossi, S. L.; Roundy, C. M.; Penate, H. N.; Kastis, M.; Schnizlein, M. K.; Le, K. C.; Haller, S. L.; Chen, R.; et al. Host oxidative folding pathways offer novel anti-chikungunya virus drug targets with broad spectrum potential. Antiviral Res 2017, 143, 246–251. DOI: 10.1016/j.antiviral.2017.04.014.

(30) Rawarak, N.; Suttitheptumrong, A.; Reamtong, O.; Boonnak, K.; Pattanakitsakul, S. N. Protein Disulfide Isomerase Inhibitor Suppresses Viral Replication and Production during Antibody-Dependent Enhancement of Dengue Virus Infection in Human Monocytic Cells. Viruses 2019, 11 (2). DOI: 10.3390/v11020155.

(31) Almasy, K. M.; Davies, J. P.; Lisy, S. M.; Tirgar, R.; Tran, S. C.; Plate, L. Small-molecule endoplasmic reticulum proteostasis regulator acts as a broad-spectrum inhibitor of dengue and Zika virus infections. Proc Natl Acad Sci U S A 2021, 118 (3). DOI: 10.1073/pnas.2012209118.

(32) Ozcelik, D.; Seto, A.; Rakic, B.; Farzam, A.; Supek, F.; Pezacki, J. P. Gene Expression Profiling of Endoplasmic Reticulum Stress in Hepatitis C Virus-Containing Cells Treated with an Inhibitor of Protein Disulfide Isomerases. ACS Omega 2018, 3 (12), 17227–17235. DOI: 10.1021/acsomega.8b02676.

(33) Kim, Y.; Chang, K. O. Protein disulfide isomerases as potential therapeutic targets for influenza A and B viruses. Virus Res 2018, 247, 26–33. DOI: 10.1016/j.virusres.2018.01.010.

(34) Santana, A. Y.; Guerrero, C. A.; Acosta, O. Implication of Hsc70, PDI and integrin alphavbeta3 involvement during entry of the murine rotavirus ECwt into small-intestinal villi of suckling mice. Arch Virol 2013, 158 (6), 1323–1336. DOI: 10.1007/s00705-013-1626-6. Aguilar-Hernandez, N.; Meyer, L.; Lopez, S.; DuBois, R. M.; Arias, C. F. Protein Disulfide Isomerase A4 Is Involved in Genome Uncoating during Human Astrovirus Cell Entry. Viruses 2020, 13 (1). DOI: 10.3390/v13010053.

(35) Varghese, F. S.; Kaukinen, P.; Gläsker, S.; Bespalov, M.; Hanski, L.; Wennerberg, K.; Kümmerer, B. M.; Ahola, T. Discovery of berberine, abamectin and ivermectin as antivirals against chikungunya and other alphaviruses. Antiviral Res. 2016, 126, 117–124. DOI: 10.1016/j.antiviral.2015.12.012.

(36) Varghese, F. S.; Thaa, B.; Amrun, S. N.; Simarmata, D.; Rausalu, K.; Nyman, T. A.; Merits, A.; McInerney, G. M.; Ng, L. F. P.; Ahola, T. The antiviral alkaloid berberine reduces chikungunya virus-induced mitogen-activated protein kinase signaling. J. Virol. 2016, 90 (21), 9743–9757. DOI: 10.1128/jvi.01382-16.

(37) Karlas, A.; Berre, S.; Couderc, T.; Varjak, M.; Braun, P.; Meyer, M.; Gangneux, N.; Karo-Astover, L.; Weege, F.; Raftery, M.; et al. A human genome-wide loss-of-function screen identifies effective chikungunya antiviral drugs. Nature Comm 2016, 7, 11320. DOI: 10.1038/ncomms11320.

